# Desmoplakin loss leads to PKC-dependent insertion of series sarcomeres and contractile dysfunction in cardiomyocytes

**DOI:** 10.1101/2025.05.15.654389

**Authors:** Ilhan Gokhan, Xia Li, Jack M. Sendek, Alex J. Mora Pagan, Fadi G. Akar, Stuart G. Campbell

## Abstract

**Background:** Mutations in *DSP*, which encodes the protein desmoplakin, lead to cardiomyopathy with unusually high penetrance. Clinical features include ventricular tachyarrhythmias, fibro-fatty infiltration of both ventricles, and ultimately dilated cardiomyopathy. While some data have been gathered to explain the electrophysiological and contractile consequences of desmoplakin cardiomyopathy, a comprehensive mechanism linking *DSP* mutations to ventricular dilation and heart failure remains elusive.

**Methods:** We use iPSC-derived engineered heart tissue (EHT) bearing a functional desmoplakin haploinsufficiency to model the heart failure phenotype that occurs in desmoplakin cardiomyopathy. Functional haploinsufficiency is secondary to a missense mutation, R451G, that results in proteolytic degradation of desmoplakin with no detectable protein. We complement functional data obtained in tissue-engineered constructs with cell biology assays in 2D cardiomyocytes to glean insights into the mechanism and mechanobiology of desmoplakin cardiomyopathy.

**Results:** Engineered heart tissues harboring a desmoplakin insufficiency recapitulate a patient phenotype notable for hypocontractility and ventricular dilation. Surprisingly, DSP-mutant tissues exhibited a shortened resting sarcomere length that was dependent on protein kinase C activity. Concurrently, mechanical load on α-catenin was increased, suggesting a mechanism by which desmosomal insufficiency redistributes force to adherens junctions. Excessive loading on adherens junctions may act as a stimulus for avid insertion of series sarcomeres, shortening the length per sarcomere, and resulting in a contractile deficit. PKC inhibition rescues shortened sarcomere length in DSP- mutant tissues, suggesting that it could be a target for future molecular therapies.

**Conclusions:** Our study uncovers a novel mechanism underlying systolic dysfunction in desmoplakin cardiomyopathy. We not only recapitulate the disease phenotype, but we identify sarcomere length regulation through altered force transmission at the intercalated disc as a previously-unrecognized mechanism.

## INTRODUCTION

Desmoplakin cardiomyopathy is a unique subtype of arrhythmogenic cardiomyopathy (ACM) with clinical features that resemble dilated cardiomyopathy^1^. ACM, which is associated with mutations in desmosomal proteins, such as *PKP2, JUP, DSC,* or *DSG*, classically presents with fibrofatty infiltration of the right ventricle and ventricular tachyarrhythmias with a high burden of sudden cardiac death^2,3^. Mutations in desmoplakin (DSP), though only accounting for ∼5% of ACM cases, cause cardiomyopathy with unusually high penetrance (57% vs 28% in *PKP2*-linked ACM), for which standard ACM risk-stratification algorithms fail^1,4^. As a result, the diagnosis of DSP-linked cardiomyopathy remains challenging, and the pathophysiological mechanisms that underly its presentation are incompletely understood^5–11^. Here, we explain how desmosomal destabilization leads to hypocontractility and dilation at the sarcomere, cell, and tissue level.

Desmoplakin is an essential component of the cardiac desmosome, connecting intermediate filaments to the intercalated disc, providing mechanical and electrical coupling between cardiomyocytes, and acting as a hub for cellular signaling^12,13^. Most disease-causing DSP variants are truncating variants that result in nonsense-mediated decay of transcript resulting in haploinsufficiency^1,4,7,14^. However, a small proportion are missense mutations that result in exposure of a cryptic calpain cleavage site resulting in a functional haploinsufficiency due to proteolytic degradation^15^. Here, we examine one such mutation, DSP^R451G^, which has been previously characterized in a cohort of symptomatic patients presenting with left ventricular failure and high burden of sudden cardiac death^16^. DSP^R451G^ is akin to desmoplakin depletion, and previous work has identified electrophysiological phenotypes that are unmasked by mechanical stretch in an *in vitro* engineered heart tissue model^17^. We build on this work to identify mechanisms of systolic dysfunction in DSP-linked ACM.

Using engineered heart tissue (EHT) expressing homozygous DSP^R451G^, we identify a hypocontractile phenotype consistent with dilation and systolic dysfunction that mimics patient presentation. Next, we subject the model to acute and chronic mechanical stretch, uncovering protein kinase C (PKC)-dependent changes in sarcomere length that account for systolic dysfunction. We show that PKC activity, which is increased in DSP^R451G^ tissues, results in sarcomere shortening and is tightly regulated by mechanical stress. Finally, we demonstrate that desmosomal destabilization results in a redistribution of force within the intercalated disc, accounting for a mechanosensitive pathway that implicates PKC-dependent assembly of series sarcomere structures in systolic dysfunction. Our work not only identifies a new mechanism for systolic heart failure in desmoplakin-linked arrhythmogenic cardiomyopathy, but also suggests novel avenues for therapy.

## METHODS

### Cell Culture and Cardiomyocyte Differentiation

Crispr-Cas9 gene editing was previously used to introduce homozygous variant R451G into the DSP gene in a healthy male iPSC line (PGP1/GM23338, Coriell Institutes for Biomedical Research)^16^. Antibiotics were not used in any cultures unless specifically noted. All cultures were regularly tested for mycoplasma by PCR. All stem cell cultures were maintained under feeder-free conditions in mTeSR1 (Stemcell Technologies). Cardiomyocytes were differentiated by biphasic Wnt modulation as previously described^60^. Briefly, iPSCs were grown to 95%+ confluence in RPMI/B27 without insulin, then treated with 20 µM CHIR99021 (day 0) for exactly 24 h to induce cardiac mesoderm. From day 3-5, cells were treated with 5 µM IWP4, and media was changed every other day thereafter. On day 9, cells were transitioned to RPMI/B27+. Between days 12-14, cardiomyocytes were metabolically purified in the presence of 4 mM lactate. After lactate selection, wells containing spontaneously-beating cardiomyocytes were trypsinized and re-plated at low density for cell expansion as previously described^61^. Cells were maintained in RPMI + 10% KOSR with 1 µM thiazovivin for 24 h, then treated with low-dose (2 µM) CHIR99021 every other day for 5 d. Expanded cardiomyocytes were reconstituted into engineered heart tissues 9-12 d after replating as described below.

### Engineered Heart Tissue (EHT) Fabrication

Expanded cardiomyocytes were seeded onto decellularized porcine LV matrix as previously described^62^. Briefly, left ventricular free wall was dissected from fresh pig hearts (J. Latella & Sons, Orange, CT) in the presence of 5x penicillin-streptomycin (P/S), cut into blocks, and cryosectioned into 150-µm slices. Slices were laser-cut into rectangular ribbons (7 x 2 mm) parallel to the fiber direction and mounted within sterilized Teflon cassettes while tension was maintained using metal pins on either end. Tissue slices were subject to hypotonic lysis (10 mM Tris-Cl, pH 7.4, 5 mM EDTA) for 3 h and decellularization (0.5% SDS in PBS) for 50 min with rotation. After extensive washing in sterile PBS, decellularized tissues were incubated overnight in DMEM + 10% FBS with 2x P/S until cell seeding.

Expanded cardiomyocytes were trypsinized 9-12 days after replating and resuspended in seeding media (high-glucose DMEM + 10% FBS with 1% NEAA, 1% glutamine, 0.18% sodium pyruvate, 1x P/S, with 10 µM Y-27632) at a density of 1,000,000 cells per EHT, containing 10% human cardiac fibroblasts (PromoCell). The following day, tissues were placed in DMEM/B27+ without antibiotics and cultured for 3 weeks before mechanical characterization.

For EHT experiments in Figure 2, expanded cardiomyocytes were seeded onto commercial decellularized scaffolds at the same density as used for standard scaffolds, but using the manufacturer’s protocol (MyoPod, Propria LLC, Branford, CT). MyoPod EHTs were cultured for 2 weeks in DMEM/B27+ and mechanically characterized using MyoLab equipment. After mechanical characterization under sterile conditions, tissues were stretched by 20% within the MyoPod cassette and returned to culture in the presence of 1x pen-strep for 48 h. After removal of antibiotics, tissues were cultured for 5 more days, then mechanically tested again.

### Mechanical Characterization of EHTs

Mechanical behavior of EHTs was characterized using a custom-built muscle fiber testing apparatus containing a KG7 force transducer (WPI) with real-time length control (ThorLabs)^62^. Tissues were unclipped from their Teflon cassettes and placed onto the force transducer using micromanipulators to avoid stretch-induced damage. Mechanical measurements were collected over a length sweep (-6% to 20% of tissue length after three rounds of preconditioning from -6% to 6%) with 1 Hz and/or 1.5 Hz pacing. At each tissue length, the EHT was equilibrated for 30 s, then 20 twitches were recorded and averaged. Tissues were perfused with CO2-equilibrated DMEM at 36 °C. Where necessary, compounds (5 µM bisindolylmaleimide VIII acetate [MedChemExpress, MCE]; 10 µM defactinib [MCE]; 1 µM Go6976 [MCE]; or 500 nM ruboxostatin HCl [MCE]) or DMSO vehicle were added to the perfusate. Analysis was performed using custom-built Matlab software. After testing, tissues were washed with PBS and flash-frozen for protein analysis or fixed overnight in 4% PFA for immunohistochemistry.

### Western Blotting

Briefly, flash-frozen EHTs were homogenized in 50-75 µL RIPA buffer with protease and phosphatase inhibitors (Thermo), rotated at 4 °C for 45 min, and the resulting lysate was clarified by centrifugation at 18,000*g*. After protein concentration determination using the BCA assay (Thermo), lysates were mixed with Laemmli buffer, boiled for 5 min, and electrophoresed on pre-cast 4-20% or AnyKD gels (Bio-Rad Mini-Protean TGX). After wet transfer to PDVF membranes, total protein was quantified using Revert Total Protein Stain (Li-Cor). Membranes were de-stained per manufacturer’s instructions, blocked for 1 h at room temperature (Li-Cor Intercept TBS Blocking Buffer), and incubated with antibodies as listed below. Signal was detected using near-IR fluorescence at 680 and 800 nm (Li-Cor Odyssey), and blots were quantified by densitometry with Image Studio (Li-Cor) software.

### Membrane Fractionation

Confluent monolayers of expanded 2D iPSC-cardiomyocytes were used for membrane fractionation experiments. Cultures were treated with either 0.5 µM mavacamten overnight, 1 µM danicamtiv overnight, 200 nM PMA for 30 min, or vehicle overnight. Cytosolic and membrane fractions were extracted using the ProteomExtract Subcellular Fractionation Kit following manufacturer’s instructions, except for the addition of phosphatase inhibitors in all extraction buffers and the use of Bradford assay (Bio-Rad) instead of BCA assay for protein quantification. Equal volumes of protein were analyzed via polyacrylamide gel electrophoresis and Western blotting as described above.

### Silver Staining

Silver staining of protein gels to analyze myosin heavy chain isoform was performed as previously described^63^. Protein extraction proceeded similar to that for Western blotting, but used a high-potassium myosin extraction buffer, containing (in mM): KCl (190), KH2PO4 (100), K2HPO4 (50), EDTA (10), Na4O7P2 (10 mM), beta-mercaptoethanol (4), pH 6.5, with 5% Triton X-100, supplemented with protease and phosphatase inhibitors. Protein concentration was calculated using the Bradford assay (Bio-Rad) and 0.5-1 µg protein was run on a 7% hand-cast polyacrylamide gel containing 10% SDS overnight at 4 mM constant current. Gels were fixed and silver-stained according to the manufacturer’s instructions (Bio-Rad Silver Stain Plus). Mouse ventricle and porcine left ventricle were used as standards for alpha- and beta-myosin heavy chain, respectively.

### Microscopy

For 2D imaging of cells for sarcomere length measurements, expanded cardiomyocytes were plated at low density on Matrigel-coated glass bottom dishes and cultured for at least 7 days in DMEM/B27+. For imaging of cell-cell junctions, expanded cardiomyocytes were seeded at higher density and allowed to grow for up to 60 d in DMEM/B27+. If required for the assay, cells were treated with drug (5 µM staurosporine for 4 h, 5 µM bisindolylmaleimide VIII acetate for 4 h, or 200 nM phorbol-myristate acetate for 30 mins) prior to fixation. Cells were fixed in 4% PFA at RT for 15 minutes, washed extensively, and permeabilized with 0.1% Triton X-100 in PBS for 15 minutes. After blocking for 1 h in 2.5% BSA in PBS, cells were treated with primary antibodies overnight at 4 °C in PBS + 0.5% BSA, followed by extensive washing and incubation with secondary antibodies at RT for 1 hr. Nuclei were counterstained with Hoechst 33528 (1:5000 in PBS).

For imaging of 3D engineered heart tissue (EHT) preparations, EHTs were washed with PBS after mechanical testing and fixed in 4% PFA overnight at 4°C. Following extensive washing, EHTs were permeabilized with 0.1% Triton X-100 in PBS for 20 minutes. For sarcomere length measurements using phalloidin-labeled actin, EHTs were treated with rhodamine-phalloidin (1:500) in 0.5% goat serum in PBS with 0.1% Triton for 2 h at RT, followed by extensive washing and counterstaining with Hoechst. For staining using antibodies, EHTs were incubated with primary antibodies in PBS + 0.5% goat serum overnight at 4°C, followed by washing, labeling with secondary antibodies and phalloidin, and counterstaining. In either case, staining was performed with the tissue pinned in its custom-made cassette to avoid shortening artifacts that could result if the tissue was released from isometric conditions. At the very last step, tissues were cut out of cassettes and mounted using ProLong Diamond antifade.

Confocal imaging was performed on a Leica Stellaris 8.

### Experimental Design and Statistical Analysis

Data were analyzed in Prism 10. For two-group comparisons between genotypes (WT and DSP-R451G), unpaired two-sided *t-*tests were used. For characterization of chronic stretch and length-dependent activation, a two-way, repeated-measures ANOVA was used, with Fisher’s LSD test for multiple-comparisons. In cases of missing data, for example due to loss of capture during pacing, a mixed-model was implemented instead of a two-way ANOVA.

### Antibodies and Labeling Reagents

| Target | Mfr./Serial Number | Application | Dilution |
| --- | --- | --- | --- |
| Desmoplakin 1/2 | Proteintech 23518-1-AP | WB | 1:1000 |
| Alpha cardiac actin | Proteintech 66125-1-Ig | WB | 1:2000 |
| Myosin heavy chain | Abcam ab50976 | WB | 1:1000 |
| PKC-alpha | Cell Signaling #59754 | WB | 1:1000 |
| PKC-epsilon | Proteintech 20877-1-AP | WB | 1:1000 |
| phospho-PKC(pan)-βII-Ser660 | Cell Signaling #9371 | WB | 1:1000 |
| phospho-PKC(pan)-γ-Thr514 | Cell Signaling #9379 | WB | 1:1000 |
| Histone H3 | Cell Signaling | WB | 1:1000 |
| N-cadherin | Cell Signaling #13116 | IF | 1:600 (cells) |
| Extended-conformation alpha-catenin (α18) | Generous gift of Dr. Akira Nagafuchi, Nara Medical University, Japan | IF | 1:600 (cells) |
| Goat anti-rat (H+L)-AlexaFluor 488 | Invitrogen #11006 | IF | 1:500 |
| Goat anti-rabbit F(ab')<sub>2</sub>-AlexaFluor 647 | Cell Signaling #4414 | IF | 1:500 |
| Goat anti-rabbit-IRDye800 | Li-Cor 926-32211 | WB | 1:10000 |
| Goat anti-mouse-IRDye680 | Li-Cor 926-32210 | WB | 1:10000 |
| Actin (rhodamine phalloidin) | Abcam ab235138 | IF | 1:500-1:5000 (cells)<br>1:250-400 (EHTs) |

## RESULTS

### DSP-R451G Results in a Hypocontractile Phenotype

Missense and truncating mutations in *DSP* have been shown to cause arrhythmogenic cardiomyopathy (ACM). In a previously-described cohort of 15 symptomatic patients harboring the DSP^R451G^ variant, 11 had biventricular or LV- predominant disease and 8 had an LVEF less than 55%^16^. The basis of DSP^R451G^ pathogenicity was previously shown to be increased susceptibility to calpain-mediated degradation, leading to a near complete loss of DSP^16^.

We previously generated engineered heart tissues (EHTs) homozygously expressing DSP^R451G^ within a normal genetic background (Fig. 1A). DSP^R451G^ EHTs displayed an almost total loss of desmoplakin protein as measured by Western blotting (Fig. 1B). We subjected DSP^R451G^ and isogenic control EHTs to a wide array of biomechanical testing. When stimulated at 1.5 Hz under isometric conditions, DSP^R451G^ EHTs displayed a hypocontractile phenotype, with peak forces that were ∼40% lower than the isogenic controls (142 µN vs 86 µN, *p* = 0.0002, Fig. 1D). This decrease in peak force was accompanied by significantly faster systolic and diastolic kinetics (time to peak, 262 ms vs 225 ms, *p* < 0.0001; time from peak to 50% relaxation, 121 ms vs 103 ms, *p* < 0.0001; Fig. 1E-G). Tissue culture and mechanical testing were conducted in DMEM, which contains physiological calcium levels (1.8 mM CaCl2). This contrasts with prior studies that used RPMI (0.42 mM Ca(NO3)2) for culture and Tyrode’s solution (1.8 mM CaCl2) for testing^16,17^. To determine if the mechanical phenotype that we uncovered in DSP^R451G^ tissues was somehow dependent on the calcium concentration, we repeated the full set of experiments using EHTs grown and tested in RPMI. Relative differences in mutant versus control tissues persisted, indicating that the observed hypocontractile phenotype is independent of extracellular calcium levels (Figure S1A-C; S2A-D). To rule out a shift in myosin heavy chain (MHC) isoform that might explain contractile differences, we ran total protein stains of the myofilament fraction. We observed that our EHT constructs express almost entirely β-MHC, the mature isoform in the human ventricle, without DSP-dependent alteration of isoform composition (Fig. 1C).

**Fig. 1:**
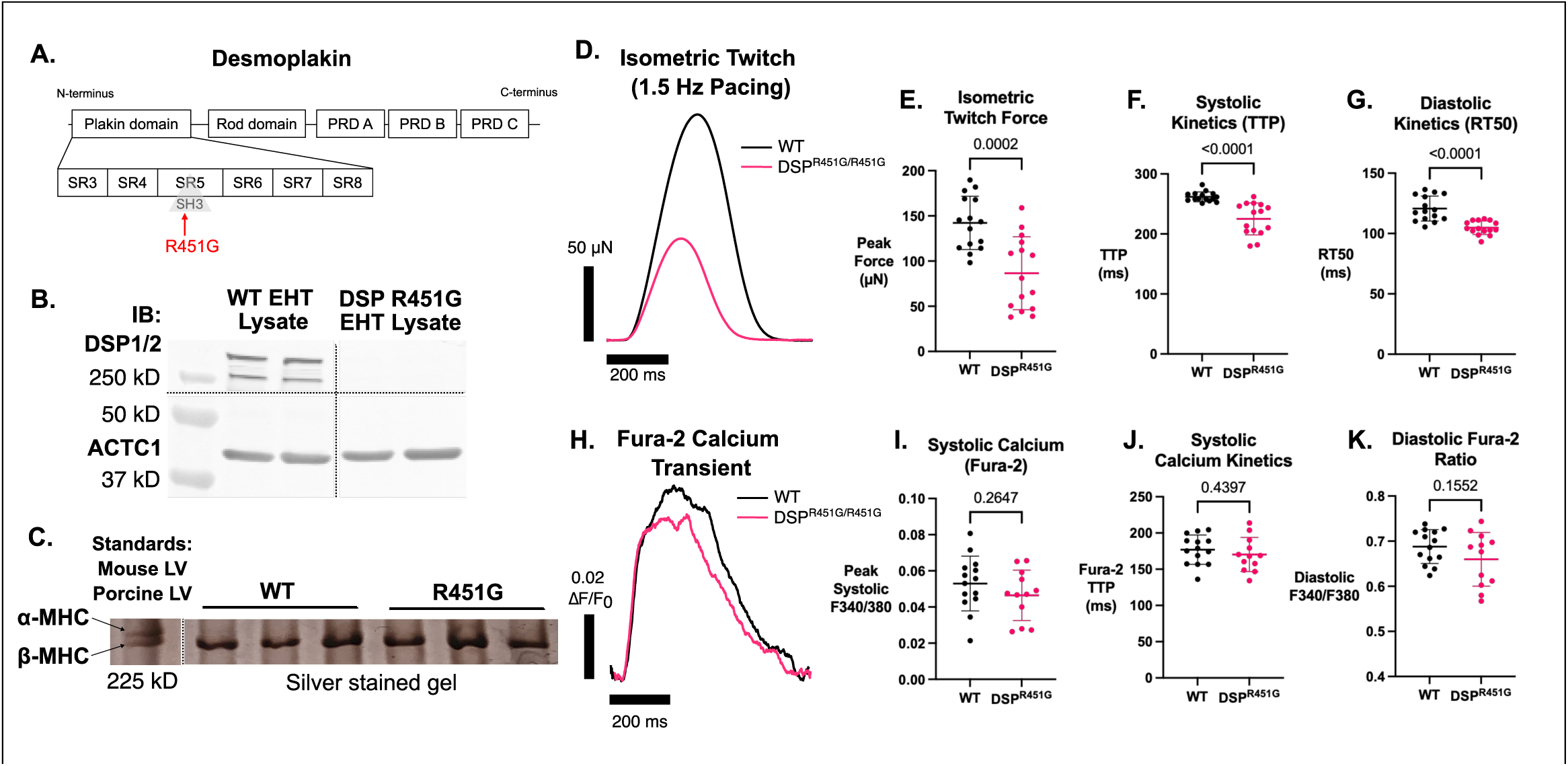
Engineered heart tissue model of mutant desmoplakin (DSP-R451G) exhibits hypocontractile phenotype**. A.** Domain diagram of desmoplakin showing location of R451G mutation within SH3 domain of plakin head. **B.** Western blot of engineered heart tissue (EHT) lysate showing total nearly loss of DSP protein in DSP^R451G^ EHTs. Dotted lines separate non-consecutive lanes from the same membrane. **C.** Silver-stained myosin gel showing exclusive expression of β-myosin heavy chain (MHC) in WT and DSP^R451G^ EHTs. Mouse ventricular and porcine left-ventricular lysates were used as a standard for α-MHC and β- MHC, respectively. **D.** Representative twitches from isometric mechanical testing of WT and DSP^R451G^ EHTs. Constructs were field-stimulated at 1.5 Hz. **E.** Isometric twitch force generated by EHTs with 1.5 Hz field stimulation (WT *N* = 15, DSP^R451G^ *N* = 15, two differentiation batches). **F.** Systolic kinetics of twitch contraction (TTP = time to peak force from stimulation time). **G.** Diastolic kinetics of twitch relaxation (RT50 = time from peak force to 50% relaxation). **H.** Representative Fura-2 calcium transients of whole tissues measured at culture length with 1.5 Hz pacing. Tissues were excited with 340 and 380 nm light and emission was collected through a GFP filter. **I.** Quantification of peak systolic calcium (WT *N* = 14, DSP^R451G^ *N* = 12); **J.** Quantification of calcium-transient upstroke (Fura-2 TTP = time to peak Fura-2- trace from stimulation); **K.** Diastolic Fura-2 ratio as measured by a ratio of emission from 340-nm and 380-nm excitation.

**Fig. 2:**
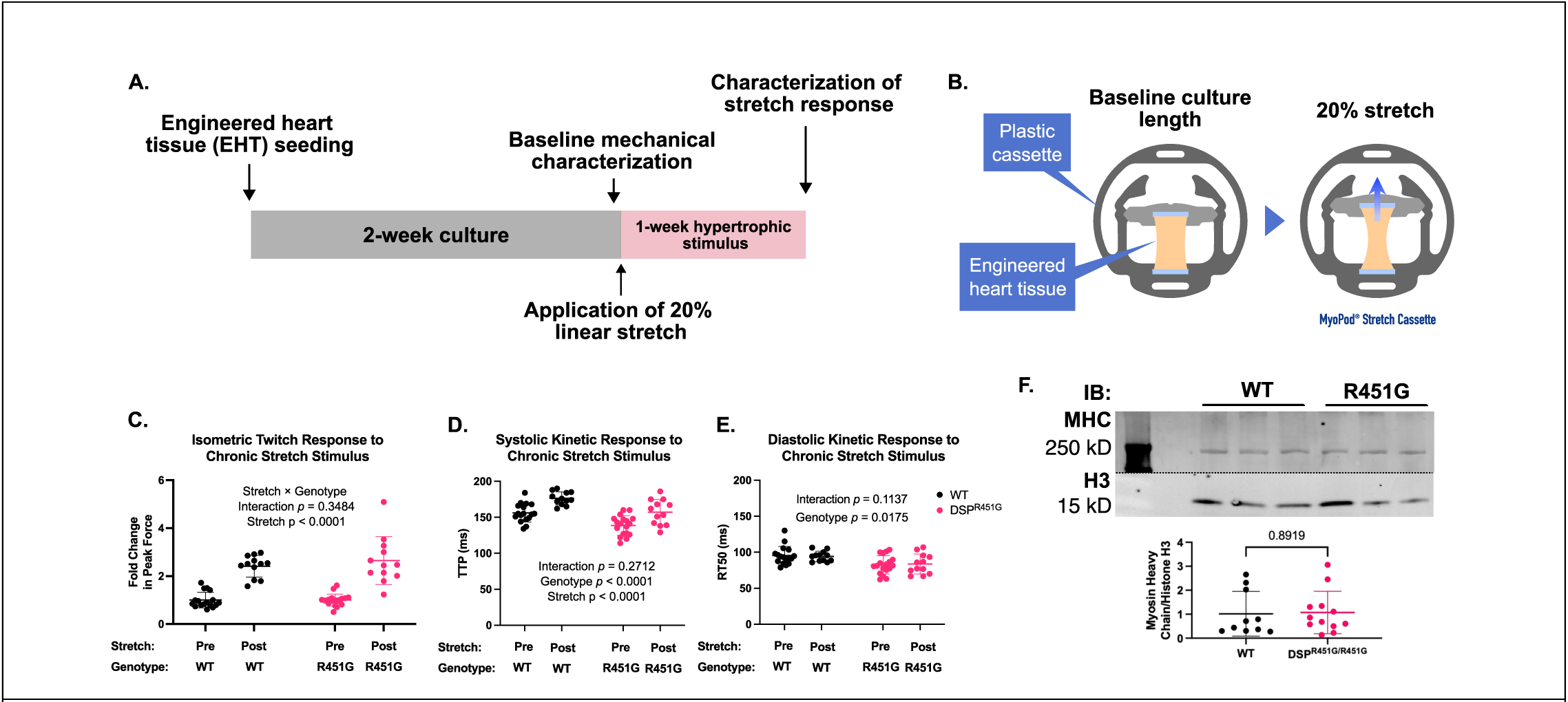
Application of chronic mechanical stress results in EHT maturation, rather than exacerbating hypocontractile ACM phenotype **A**. Diagram of experimental design, consisting of 2 weeks of culture after tissue seeding, baseline characterization, and 1 week of culture with 20% biomechanical stretch. **B.** Schematic of tissue cassette that enables tissue stretch. A 20% stretch stimulus is applied at once after baseline mechanical testing. **C.** Fold-change in isometric twitch force before and after 20% mechanical stretch. WT and DSP^R451G^ EHTs responded equally to this hypertrophic stimulus. WT, pre-stretch N = 17, post-stretch N = 12; DSP^R451G,^ pre-stretch N = 18, post-stretch N = 12, three differentiations per genotype. Statistical design used two-way ANOVA with repeated measures to account for paired comparison. **D.** Systolic kinetics (time from stimulation to peak force, TTP) were not affected by mechanical stretch in a genotype-specific manner. **E.** Diastolic kinetics (time from peak force to 50% relaxation) were not affected by mechanical stretch in a genotype-specific manner. **F.** Western blot showing equal protein expression of myosin heavy chain (MHC) in WT and DSP^R451G^ EHTs after stretch (WT N = 10, DSP^R451G^ N = 12, unpaired Student’s *t*-test.)

Because calcium handling is frequently dysregulated in ACM models, including those involving PKP2 deficiency^18,19^, we next assessed calcium transients in WT and DSP^R451G^ tissues using Fura-2 imaging of intact EHTs. Remarkably, we found no differences in the shape and amplitude of the integrated calcium transient (Fig. 1H-K). Although there were subtle kinetic differences in acute stretch-induced remodeling of the calcium transient (Fig. S3A-C), there were no changes in systolic peak calcium at culture length or under acute stretch. As a result, the hypocontractile phenotype of DSP^R451G^ cannot be explained by changes in calcium handling. We observed arrhythmic behavior in the form of faster spontaneous beating that caused loss of capture in approximately half of DSP^R451G^ EHTs at 20% acute stretch. However, diastolic calcium ratios were the same across genotypes (Fig. S3D). None of the measured tissue-level calcium transient parameters showed sufficient dysregulation to suggest the presence of an arrhythmogenic substrate or trigger. These findings are consistent with our prior work demonstrating that unmasking major phenotypes such as reductions in Cx43 expression and conduction velocity require dynamic culture conditions, an approach not employed in the current study^17^.

### Chronic Mechanical Stretch Matures WT and DSP-R451G EHTs Equally

Previous work has demonstrated that dynamic culture is necessary to produce a disease phenotype in another DSP-linked ACM model^20^. Accordingly, we challenged WT and DSP^R451G^ EHTs with chronic mechanical stretch to determine whether this mechanical perturbation would exacerbate the basal disease phenotype. Using a paired experimental design, we mechanically tested EHTs before and after one week of 20% linear stretch (Fig. 2A-B). Baseline data recapitulated the hypocontractile phenotype previously shown (Fig. S4). Remarkably, both WT and DSP^R451G^ produced two-fold higher contractile force after chronic stretch, without a genotype-dependent interaction (Fig. 2C). Systolic kinetics were slightly slower in both genotypes after stretch, and diastolic kinetics remained unchanged by stretch, with insignificant stretch-genotype interactions. The lack of genotype-specific changes in kinetic behavior suggests that both groups underwent appropriate physiological remodeling after the onset of a pro-hypertrophic stimulus.

We expected that a chronic mechanical stress would further worsen contractility in DSP^R451G^ tissues, presumably because of desmosomal destabilization secondary to DSP loss. Instead, DSP-mutant tissues responded just as well to a chronic-stretch stimulus as their WT counterparts. Because the major endpoint of stretch-induced force upregulation is insertion of parallel sarcomeres, we surmise that these mechanisms remain intact in our DSP-depletion model^21–23^. Indeed, protein expression of myosin heavy chain (MHC), normalized to histone H3 expression, was equal in WT and DSP^R451G^ EHTs after stretch (Fig. 2F). Seeing that DSP^R451G^ tissues respond robustly to chronic stretch suggests that factors beyond simple destabilization of cell-to-cell desmosomal contacts is contributing to the baseline hypocontractile phenotype.

Accordingly, we next applied acute stretch to WT and DSP-mutant EHTs to determine whether more subtle alterations in contractility could be unmasked.

### Sarcomere Length Differences Explain Enhanced Length-Dependent Activation in DSP- R451

We next characterized the acute response of DSP^R451G^ EHTs to stretch. Length- dependent activation, a sarcomere-level corollary of the Frank-Starling effect, reflects sarcomere overlap and calcium sensitivity of myofilaments^24^. We performed acute length sweeps from -6% to +20% of culture length while recording twitch contractions at 1.5 Hz stimulus frequency (Fig. 3A). As expected, peak force production by DSP^R451G^ was lower than WT at every length (Fig. 3C). However, when we normalized peak force by culture-length force (i.e. at 0% stretch), DSP^R451G^ EHTs slightly outperformed their WT counterparts (Fig. 3D, *p*-value of genotype-stretch interaction 0.0092). In other words, DSP^R451G^ tissues were able to produce a greater fold-change increase in force when stretched, despite having a contractile deficit at baseline. Such a surprising finding prompted us to examine sarcomere-level factors that may explain the baseline phenotype of DSP^R451G^ as well as its acute-stretch response.

**Fig 3:**
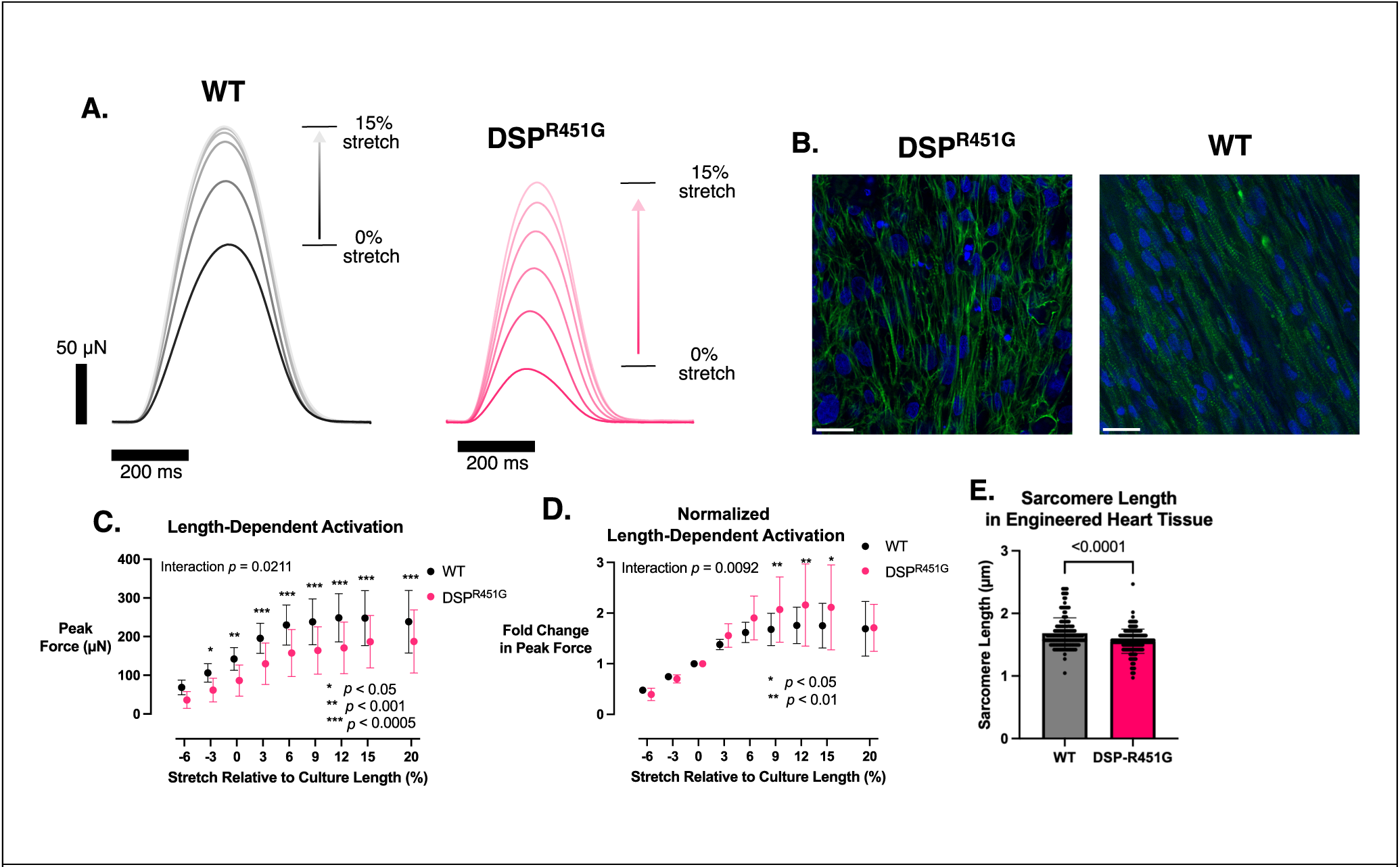
DSP^R451G^ EHTs exhibit enhanced length-dependent activation, secondary to shorter diastolic sarcomere length. **A.** Representative isometric twitches at 1.5 Hz pacing showing lower baseline force production in DSP^R451G^ EHTs, with a greater fold-change in force production over a length sweep from 0 to 15% stretch (3% intervals). **B.** Whole-mount confocal micrographs of fixed, phalloidin-stained EHTs, showing aligned sarcomeres. Scale bar = 20 µm. **C.** Length- dependent activation curve demonstrating that both WT and DSP^R451G^ tissues display Frank-Starling behavior. Two-way repeated-measures ANOVA used for statistical analysis (WT and DSP^R451G^ *N* = 15) **D.** Normalization of Frank-Starling curve in Panel C suggests that DSP^R451G^ tissues display enhanced length-dependent activation. Peak force values were normalized to culture length (i.e. 0% stretch). **E.** Sarcomere length difference between WT (N = 3) and DSP^R451G^ (N = 4) EHTs. From each blinded confocal z-stack, 5-10 myofibrils were traced as line scans and peak-to-peak distance calculated in Matlab. Total number of sarcomeres analyzed: WT *n* = 180, DSP^R451G^ *n* = 161. Each biological replicate was from a separate differentiation.

The length-dependent activation curves of WT and DSP^R451G^ tissues appear horizontally superimposable (Fig. 3C), suggesting that a difference in sarcomere length may underly contractile deficits in DSP^R451G^ tissues. According to the classic force-length relationship in striated muscle, maximal force generation occurs at an optimal sarcomere length that ensures ideal actin-myosin filament overlap^25^. When sarcomeres are overstretched (i.e. the Z-discs are too far apart), filament overlap decreases, reducing actomyosin cross-bridge formation. When sarcomeres are overly compressed, myosin heads may be sterically hindered by Z-disks or disrupted by overlapping thin filaments from adjacent half-sarcomeres, also limiting force production. We measured diastolic sarcomere lengths in WT and DSP^R451G^ EHTs using confocal microscopy and found that sarcomere length was shorter in DSP-mutant EHTs (1.68 µm vs 1.55 µm, *p* < 0.0001, Fig. 3B, E). Moreover, the relative change in sarcomere length (∼8%) corresponded to the horizontal shift between the WT and DSP^R451G^ length-dependent activation curves. Consequently, a shortened baseline sarcomere length may explain much of the hypocontractile phenotype seen in the DSP^R451G^ background, as well as the potential for enhanced length-dependent activation. Though sarcomere disarray is a well-appreciated phenotype in ACM, this is the first demonstration, to our knowledge, that implicates shorter sarcomeres as a potential cause of hypocontractility, and thereby dilation, in ACM^26^.

### Protein Kinase C Regulates Sarcomere Length in WT and DSP-R451G Cells

To verify DSP-dependent changes in sarcomere length observed in 3D tissues, we repeated sarcomere length measurements in a simpler system consisting of expanded iPSC-derived cardiomyocytes plated on Matrigel (Fig. 4A, top). Recapitulating results obtained in fixed EHTs, DSP^R451G^ cells had shorter sarcomere lengths than isogenic controls (1.96 µm vs 1.79 µm, *p* < 0.0001, Fig. 4B). We found that sarcomere lengths of plated cardiomyocytes were slightly longer than those in engineered heart tissue, but this may result from a relaxation artifact that occurs when EHTs are removed from isometric cassettes and mounted on slides during the last step of imaging.

**Figure 4:**
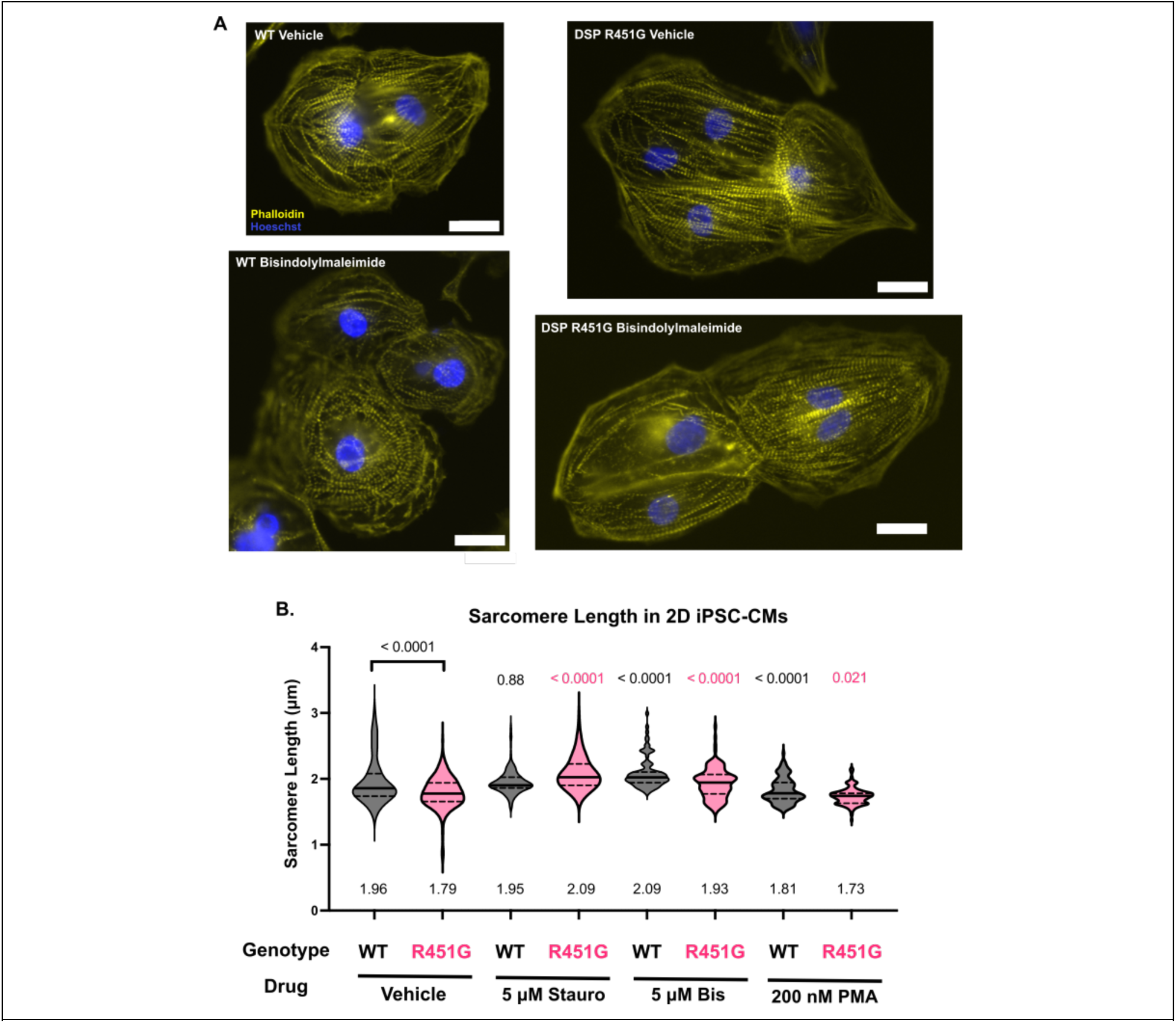
PKC inhibition rescues sarcomere length deficit in 2D iPSC-derived cardiomyocytes **A.** Representative micrographs of cell clusters stained with phalloidin and Hoechst. Cells were treated with 0.05% vehicle control or 5 µM pan-PKC inhibitor bisindolylmaleimide VIII acetate for 4 h prior to fixation and labeling. **B.** Drug screen of PKC-activating and inhibiting compounds. Stauro = staurosporine; Bis = bisindolylmaleimide VIII; PMA = phorbol myristate acetate. Cells were treated with indicated compound for 4 h prior to fixation and immunofluorescence. Numbers below each violin are average sarcomere length for each condition. Statistics used two-way ANOVA. Numbers above each violin are *p* value for multiple comparisons between drug treatment and vehicle control of same genotype. Interaction *p-*value for drug treatment is <0.0001. Each measurement consists of at least seven fields of view containing at least 25 cardiomyocytes, from which line scans of 5-10 sarcomeres were generated to calculate sarcomere length.

Regardless, the relative ∼8% decrease in sarcomere length was consistently observed, whether 3D EHTs or 2D-iPSC cardiomyocytes were used.

Previous literature has implicated several kinases in sarcomere length regulation including focal adhesion kinase (FAK) and protein kinase C (PKC)^27^. Accordingly, we manipulated these pathways in 2D culture to see if we could rescue the sarcomere length deficit in DSP^R451G^. While DSP^R451G^ cardiomyocytes had enhanced staining of phospho-T397-FAK, pharmacological blockade with defactinib only modestly reduced staining, and had no effect on 3D EHT contractility (Fig. S6, S7). However, acute treatment of DSP^R451G^ cardiomyocytes with staurosporine, a potent but nonspecific PKC inhibitor, rescued the sarcomere length deficit and lengthened sarcomeres beyond the WT value, from 1.79 µm to 2.09 µm (Fig. 4B). To avoid potential off-target effects of staurosporine, we repeated the treatment with bisindolylmaleimide VIII acetate, a more specific, pan-PKC inhibitor. Similarly, we found that sarcomere length was rescued in DSP^R451G^ cells, from 1.79 µm to 1.93 µm. Given the multifaceted role of the PKC pathway in cardiomyocyte physiology, we sought to validate the sarcomere length effect using a PKC activator, phorbol 12-myristate 13-acetate (PMA). Acute treatment with 200 nM PMA significantly decreased sarcomere length in both WT and DSP^R451G^ cells, with a greater decrease in WT cells (1.96 to 1.81 µm [WT], vs 1.79 to 1.73 µm [DSP]). These findings—particularly reduced sarcomere length following treatment with a PKC agonist known to promote sarcomere assembly^28^—suggest that excessive addition of sarcomeres in series can shorten the length of an individual sarcomere. Series sarcomere insertion into myofibrils with maintained cell length would result in a small length per sarcomere, analogous to a “compression” effect. As a result, DSP^R451G^ cells and tissues operate on a suboptimal region of the muscle length–tension curve, which may explain both reduced peak force production at culture length and potentiated length-dependent activation.

### PKC Inhibition Rescues Sarcomere Length in Engineered Heart Tissue with Genotype-Specific Effects on Function

We next considered whether PKC inhibition would rescue sarcomere length in DSP^R451G^ engineered heart tissue, a more physiologically-relevant model compared to 2D cardiomyocytes. Rather than use staurosporine, which has many off-target effects, or bisindolylmaleimide, which decreased contractility in WT EHTs (Fig. S5A), we turned to ruboxistaurin, a PKC inhibitor that showed benefit in murine models of dilated cardiomyopathy and was well-tolerated in clinical trials^29–31^. With 24 h of treatment, ruboxistaurin restored sarcomere length in DSP^R451G^ (WT vehicle length, 1.87 µm; vehicle DSP^R451G^, 1.76 µm; ruboxistaurin-treated DSP^R451G^ 1.94 µm, interaction *p-*value 0.016; Fig. 5A). This suggests that the PKC-dependance of sarcomere length in DSP^R451G^ cells persists whether they are grown on flat substrates or within a 3D tissue.

**Fig. 5:**
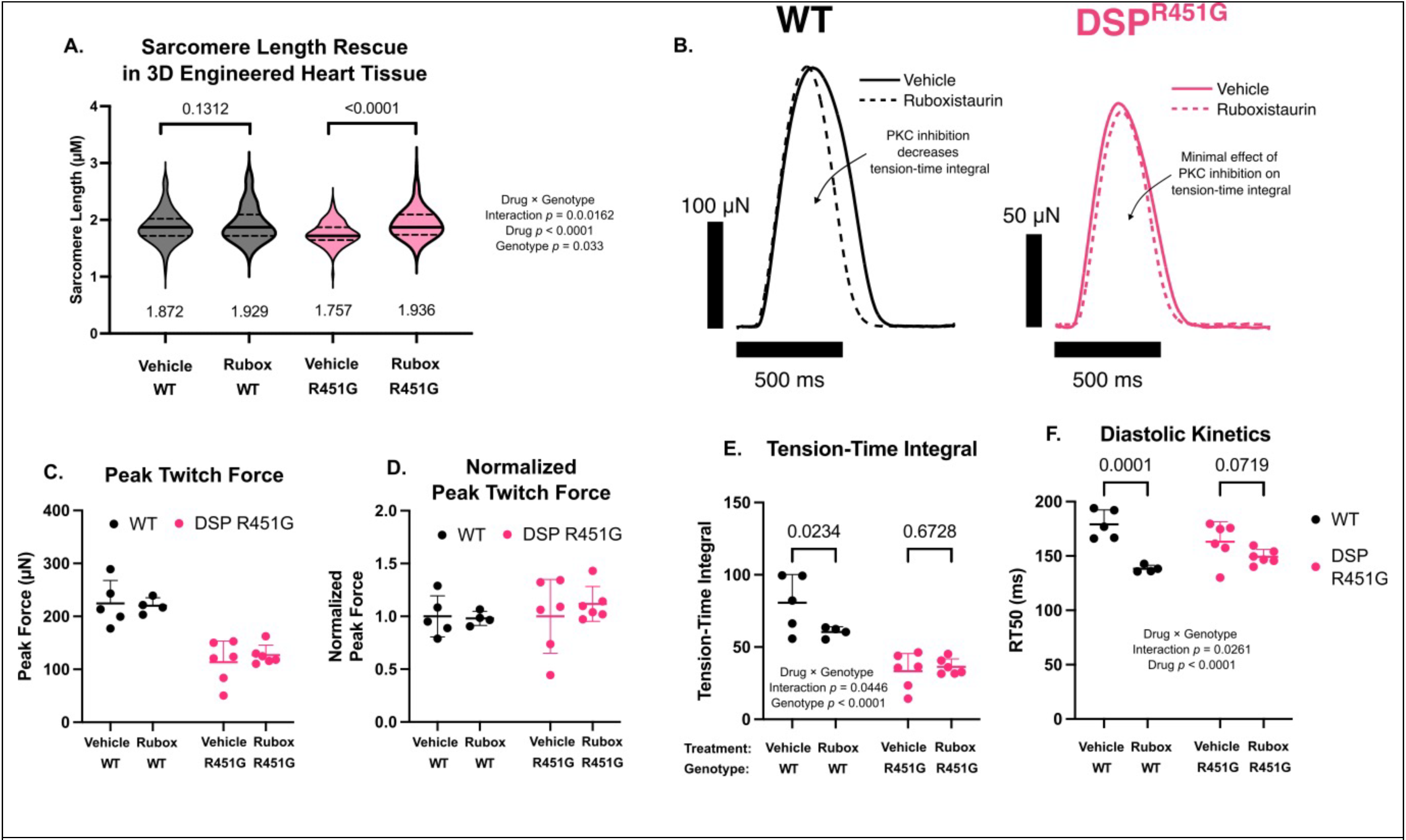
Ruboxistaurin rescues sarcomere length in 3D EHTs, with genotype-specific effects on contractile function. **A.** Ruboxistaurin rescues sarcomere length in DSP^R451G^ EHTs (two-way ANOVA drug-genotype interaction *p* = 0.0162, N = 3 EHTs per group, with *n* = 88-136 sarcomeres per condition). **B.** Representative twitches showing how ruboxistaurin decreases tension-time integral in WT tissues but has a neutral effect in DSP^R451G^ EHTs. **C.** Isometric twitch force at 1 Hz pacing of vehicle- and ruboxistaurin-treated EHTs. WT vehicle, N = 5; WT ruboxistaurin, N = 4; DSP^R451G^ vehicle, N = 5; DSP^R451G^ ruboxistaurin, N = 6; two-way ANOVA used for statistical comparison. **D.** Twitch force normalized to each genotype’s vehicle. **E.** Tension-time integral (area under twitch curve) of WT and DSP^R451G^ EHTs, showing a genotype-specific effect of ruboxistaurin, driven mostly by relaxation kinetics. **F.** Diastolic kinetics (RT50, time from peak force to 50% relaxation) in DSP^R451G^ and WT tissues.

We also performed functional assays on ruboxistaurin-treated DSP^R451G^ tissues and found genotype-specific effects on many contractile parameters. Ruboxistaurin decreased the tension-time integral (TTI) in WT tissues but had no effect in DSP^R451G^ tissues (Fig. 5E). These changes were driven by a large decrease in relaxation time in WT tissues (179 vs 138 ms, *p* = 0.0001), and a statistically-insignificant change in DSP^R451G^ tissues (163 vs. 149 ms, *p* = 0.072, Fig. 5F). Ruboxistaurin also resulted in a ∼12% increase in force compared to vehicle in DSP^R451G^ EHTs, though the effect was not statistically significant and did not amount to a rescue relative to WT (Fig. 5C-D). Titration of drug dosing may identify an optimal dose at which beneficial changes in DSP^R451G^ sarcomere length overcome any off-target effects. We next sought to explain why and how mutant desmoplakin could lead to enhanced PKC signaling, a phenomenon that has not previously been reported in the literature.

### Altered Mechanical Homeostasis Explains Enhanced Baseline Activation of Protein Kinase C in DSP-Mutant Cells

We have shown that sarcomere length is shortened in DSP^R451G^ cardiomyocytes and tissues, and that pharmacologic PKC blockade with three different compounds rescues the phenotype in 2D and 3D systems. PKC activation, conversely, shortens sarcomere length (Fig. 4B), as previously described^27^. This suggests that basal PKC activity in DSP^R451G^ cells may be enhanced. Using an antibody against the hydrophobic- tail phospho-site of PKCβII (Ser660) that marks a conserved activation mechanism of all PKC enzymes^32^, we showed that pan-PKC activity is enhanced in DSP^R451G^ tissues vs WT tissues (Fig. 6A-C). This baseline increase in activity is consistent with excessive insertion of series sarcomeres resulting in a shorter length-per-sarcomere. However, in tissues that were subject to 20% stretch for 1 week, PKC phosphorylation was equal in DSP^R451G^ vs WT tissues. These results corroborate previous literature demonstrating stretch-induced activation of PKC, but also suggest a ceiling-level effect in DSP-mutant tissues, wherein stretch does not further activate PKC^33^. In other words, stretch-sensitivity of PKC activation is lost. While many factors may cause enhanced PKC phosphorylation in DSP^R451G^ EHTs, we hypothesize that desmosomal disruption alters mechanical homeostasis in a way that promotes stretch-activated PKC activation at baseline. To investigate this, we turned to a 2D cardiomyocyte immunofluorescence assay to stain intercalated disc and adherens junction proteins.

**Fig. 6:**
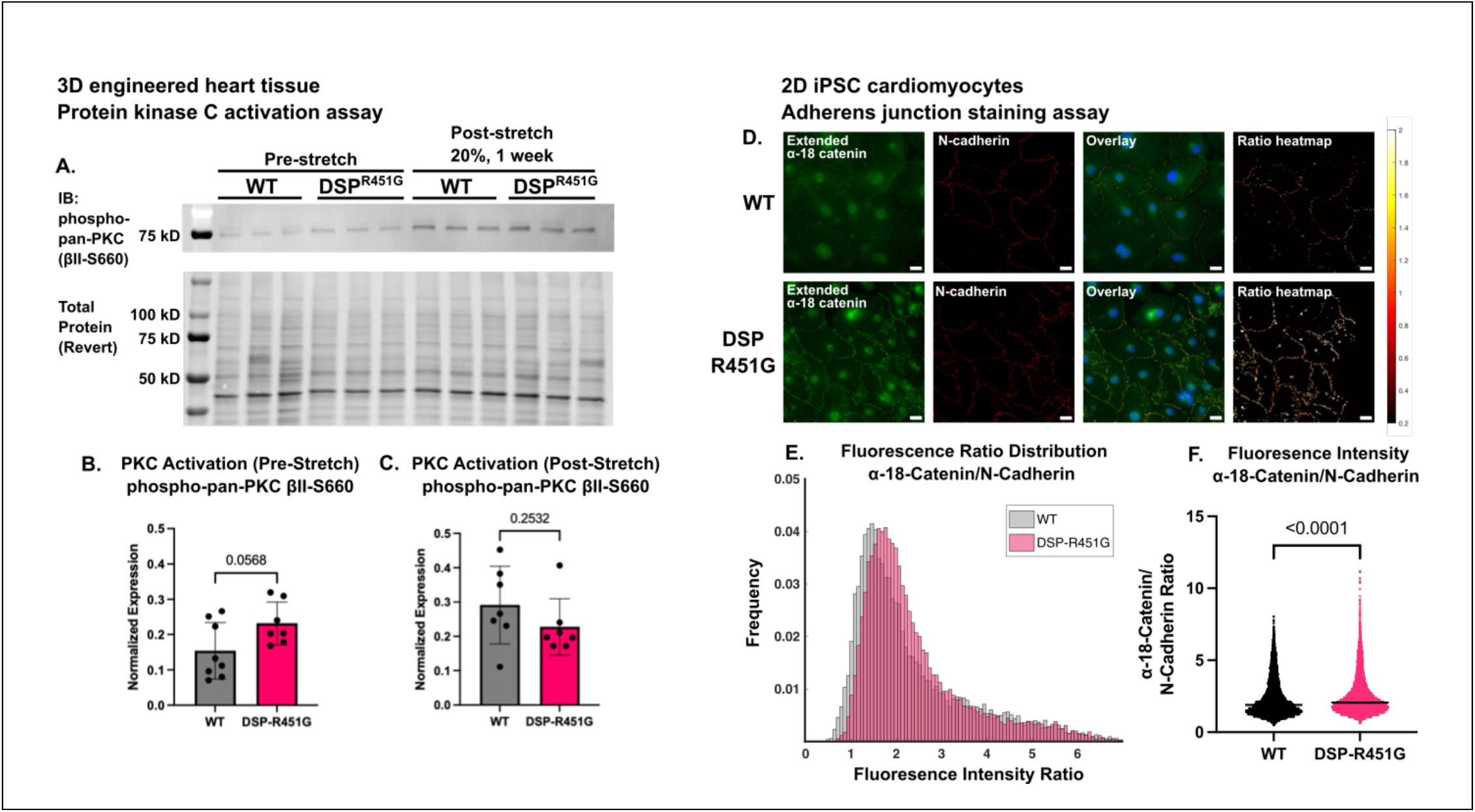
DSPR451G EHTs have enhanced PKC phosphorylation and a loss of stretch-sensitive PKC activation, while DSP^R451G^ myocytes exhibit increased mechanical loading at cell-cell junctions **A.** Western blot of EHT lysate probed for active (phosphorylated) pan-PKC. βII-S660 site corresponds to the hydrophobic tail site of activated PKC. **B.** and **C.** Quantification of phospho-PKC normalized to total protein (Li-Cor Revert). EHTs in stretched and non-stretched cassettes were from different experiments so were not analyzed together. Statistical comparison made using Student’s *t*-tests. Pre-stretch samples came from 3 (WT) and 4 (DSP) differentiations, post-stretch samples came from 3 differentiations (WT and DSP). **D.** Widefield immunofluorescence staining of 6-week-old, expanded 2D iPSC-derived cardiomyocytes. Extended-conformation α18 corresponds to an epitope of α-catenin unmasked by mechanical stress. N-cadherin staining used as fiduciary adherens junction marker. Images were processed using watershed algorithm to identify junctions. Scale bar, 20 µm. **E.** Histogram of α18-catenin to N-cadherin fluorescence ratio at segmented junctions. **F.** Quantification of data in **E**., with Student’s *t*-test for comparison. Data obtained from two independent differentiations of each cell line, with *N* = 27 monolayer fields-of-view, 10844 segmented junctions (WT) and *N* = 33 monolayer FOVs, 18869 segmented junctions.

Within the intercalated disc, desmosomes are located in close proximity to adherens junctions (AJ), which connect to force-generating actomyosin filaments through a complex network of proteins including vinculin, α- and β-catenin and plakoglobin. Inspired by the previous finding that plakophilin loss in a single-cell context results in maintained or increased traction force, we hypothesized that disruption of the desmosome would subject greater force on the cytoplasmic face of neighboring adherens junctions^26,34^. Using an antibody that recognizes an extended conformation of α-catenin under mechanical load (α18), we performed immunostaining in 2D cardiomyocyte monolayers, using N-cadherin as a fiducial adherens junction marker^35,36^ (Fig. 6D). N-cadherin expression and localization in ARVC biopsies is largely indistinguishable from control samples^37^. After segmenting adherens junctions using the N-cadherin channel and normalizing by N-cadherin intensity, we found enhanced junctional α18-catenin staining in DSP^R451G^ cells. Though α18-catenin is only a proxy for the mechanical state of AJs and not a direct force readout, it suggests that mechanical force that would partially be borne by desmosomes is instead transmitted to AJs in the setting of desmoplakin knockout. While the nature of force redistribution in this assay is different from stretch-activated PKC activation described previously, taken together, these experiments may suggest that load redistribution from desmosomes to AJs may activate PKC in DSP^R451G^ constructs. Consequently, this provides a mechanosensitive mechanism for PKC-dependent sarcomere addition in a hypocontractile model of DSP- linked arrhythmogenic cardiomyopathy.

## DISCUSSION

Arrhythmogenic cardiomyopathy is often associated with systolic heart failure in the setting of desmoplakin mutations. Using 3D engineered heart tissue harboring desmoplakin haploinsufficiency, we recapitulate this hypocontractile phenotype and identify shortened sarcomere length as a contributing pathogenic mechanism.

Corroborating previous literature on series sarcomerogenesis, we show that PKC activity directly modulates average sarcomere length^27^. Simultaneously, we measure increased mechanically-loaded α-catenin at junctions between DSP^R451G^ cells, suggesting a redistribution of force from desmosomes to adherens junctions. As a result, adherens junctions, which are already under mechanical stress because of their connection to cycling actomyosin crossbridges, undergo excessive loading, which is a well-appreciated stimulus for sarcomerogenesis^38^. Therefore, we conclude that avid series sarcomere insertion may be a mechanism that gives rise to ventricular dilation in cases of desmoplakin haploinsufficiency and possibly in ACM in general (Fig. 7).

**Fig. 7:**
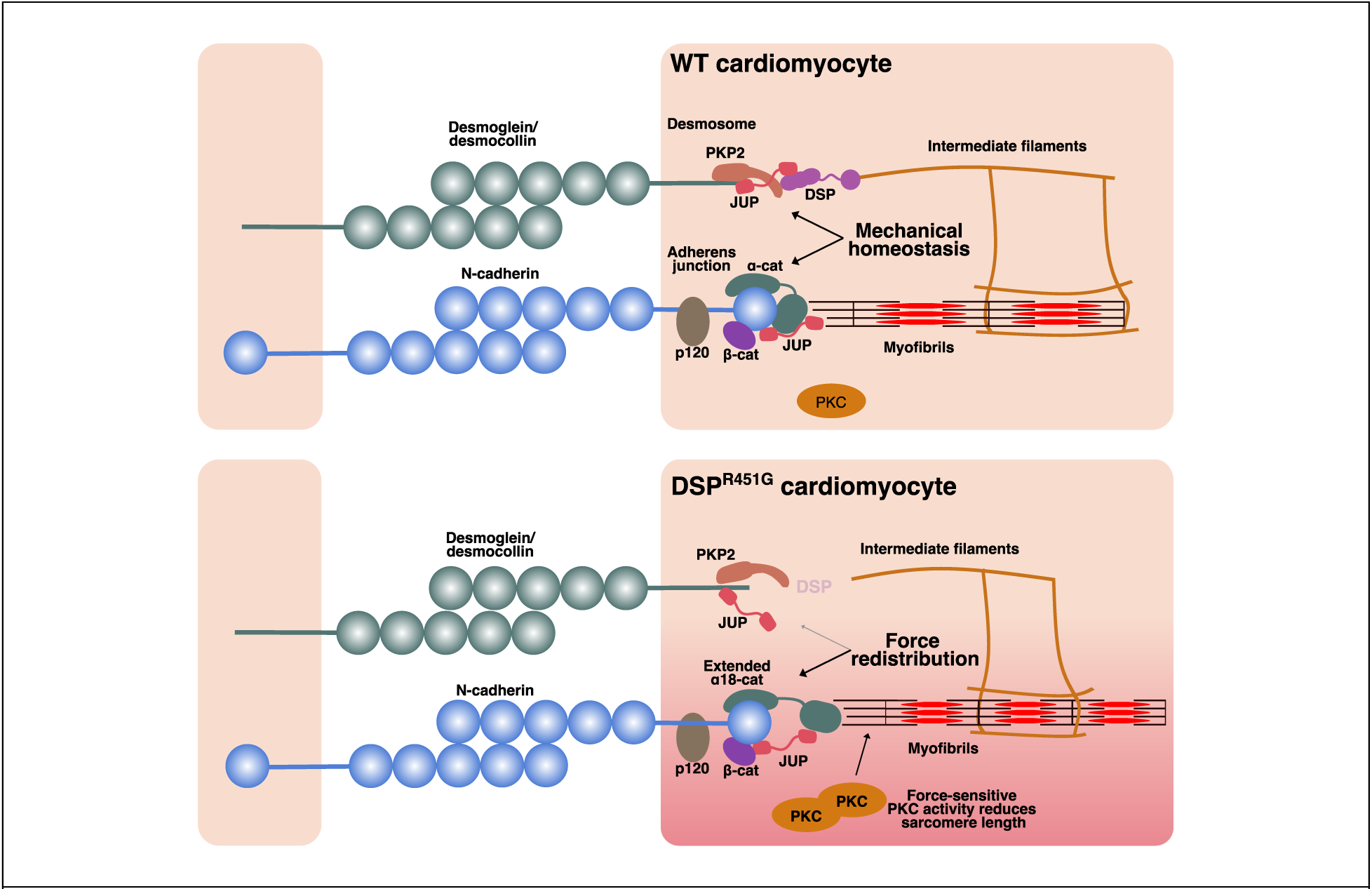
Working model of force redistribution between desmosomes and adherens junctions resulting in PKC-mediated insertion of series sarcomeres in the setting of DSP haploinsufficiency.

Our findings are broadly consistent with previous *in vitro* and *in vivo* models of ACM. Overexpression of mutant desmoplakin (R2843H) has been shown to cause ventricular dilation and damage to intercalated disc ultrastructure, while heterozygous *Dsp* knockout mice develop fibrofatty infiltration through nuclear localization of plakoglobin and reduction of Wnt/beta-catenin signaling^14,39^. Meanwhile, a desmin- knockout mouse, though not a model of ACM, produces less active force with maintained calcium sensitivity, suggesting a key role for intermediate filaments in active contraction^40^. Though we did not examine intermediate filaments in our study, mutant DSP^R451G^ would likely have a similar effect by removing the anchoring point for desmin filaments. Finally, a heterozygous PKP2 haploinsufficient mouse exhibited decreased total actin expression with aging, resulting in decreased fractional and sarcomere shortening, though this finding was limited to the RV and not associated with a change in sarcomere length^41^.

The role of the sarcomere in ACM disease progression has only recently been investigated. Elegant work in an iPSC-based PKP2-truncation model has shown that increased turnover of junctional N-cadherin results in global sarcomere disarray, though sarcomere length was not reported^26^. These findings appear to contradict our results, which show increased mechanically-loaded α-catenin at junctions. However, this PKP2- truncation model only exhibits a small-to-moderate decrease in DSP expression, with desmosomes that may be more intact than those in our model, in which DSP is entirely absent. As a result, we would not expect this model to exhibit a force redistribution from desmosomes to adherens junctions resulting in the sarcomere length-phenotype we observed. Likewise, an atrial cell-culture model of PKP2 deletion also showed sarcomere disarray, but sarcomere length was normal in the 25% of mutant cells that contained sarcomeres^42^. In contrast, a mouse model of plakoglobin (*JUP*) deletion more closely resembles our findings. In this system, desmosomes are entirely absent but adherens junctions are intact, and markers of ventricular wall stress, such as BNP, were dramatically increased^43^. Despite differences in the species and model system, this is an organ-level corollary of our finding that mechanical loading at adherens junctions increases when desmoplakin is lost. Taken together, the mechanism of hypocontractility in arrhythmogenic cardiomyopathy may depend on which individual protein of a desmosome is disrupted, because each component contributes to mechanical homeostasis or scaffolding of signaling proteins in a slightly different way.

Beyond implicating sarcomere length as a key contributor to ACM pathogenesis, our findings also unveil a previously unrecognized role for protein kinase C (PKC) signaling in this disease. To our knowledge, PKC has not been previously linked to ACM. However, its broad influence on cardiac physiology and remodeling is well established, and notably, PKP2, a core desmosomal protein implicated in ACM, has been shown to scaffold PKCα in the skin. These connections raise the possibility that aberrant PKC signaling may contribute to cardiomyocyte dysfunction in ACM^44–46^.

PKCα, which is the predominant isoform in the heart, phosphorylates many cardiac-specific substrates including the PEVK motif of titin, troponin I, troponin T, and phospholamban^47–49^. PKCα-KO mice display enhanced contractility with adaptive response to pressure overload, while PKCα overexpression results in heart failure^50^. Notably, PKCα localizes to the intercalated disc in a genetic model of DCM, and its inhibition by both endogenous proteins and drugs is therapeutic^29,51^. Though treatment with PKC inhibitor ruboxistaurin did not enhance contractility in DSP^R451G^ tissues, it exhibited a genotype-specific effect on tension-time integral and successfully rescued sarcomere length. This confirms PKC activity as a previously-unrecognized mechanism in ACM and may explain overlapping presentations between DSP-linked arrhythmogenic cardiomyopathy and dilated cardiomyopathy. Notably, PKC inhibition has shown beneficial effects in other models of cardiovascular disease, and ruboxistaurin, in particular, was well-tolerated in Phase I-III clinical trials for other indications^30,31,52^.

We cannot rule out contributions from other PKC isoforms to our results since we used a pan-PKC phospho-antibody in our PKC activation assay. Of particular interest is PKCε, which phosphorylates focal adhesion kinase (FAK), thereby promoting longitudinal sarcomere growth from costameres^53,54^. Indeed, we found increased phospho-T397-FAK staining in DSP^R451G^ cells (Fig. S6); however, acute treatment with a phospho-FAK inhibitor (GSK-2256089) did not rescue sarcomere length, and chronic treatment with another inhibitor, defactinib, did not rescue tissue hypocontractility. It is possible that other phospho-sites may be involved, and that 2D culture poses physiologic limitations to studying this pathway^55^. And although mechanotransduction at focal adhesions and costameres is an essential process for sarcomere assembly^27,38,56^, we focused on the intercalated disc and adherens junction as the site for series sarcomere assembly in DSP-linked ACM^57^. Notably, the addition of parallel sarcomeres, which is a classic finding in pressure overload-induced hypertrophy, was unlikely to have been affected by genotype in our system, because application of a chronic stress stimulus (20% stretch for 1 week) resulted in identical contractile remodeling and force enhancement in WT and DSP-mutant EHTs.

We acknowledge that *in vitro* disease modeling of ACM is exquisitely sensitive to the nature of the engineered tissue construct. Our study employed engineered heart tissue that was grown under isometric conditions, rather than with dynamic culture, which may have resulted in a less-realistic disease phenotype^17,20^. Also, the sarcomere-length phenotype that we observe may not directly translate *in vivo,* because the heart does not contract under isometric conditions, and other mechanisms may compensate for a sarcomere-length deficit. Regarding limitations of our model system, cardiomyocytes that have been differentiated from iPSCs are known to more closely resemble fetal-like myocytes rather than adult tissue^58,59^. Though iPSC-derived cardiomyocytes contain relatively immature contractile units, myofibrils in our 3D constructs are relatively well-aligned, even in the DSP mutant (Fig. 3B) and sarcomere lengths in our constructs (Fig. 4, 5) are relatively consistent with physiological values. However, our 2D iPSC-derived cardiomyocytes are cultured in a less physiologic context than our 3D constructs, and display size and morphology that are both smaller and less anisotropic than native cardiomyocytes *in vivo*. As a result, it may be difficult to delineate the precise subcellular localizations from which series sarcomeres are added. Additionally, because these monolayers were cultured on glass surfaces atop a Matrigel mattress, they may be subject to unrealistic mechanical loads that do not fully recapitulate cell-matrix interactions within native myocardium. Nevertheless, these limitations would be expected to affect each genotype the same way.

In summary, our study uncovers a novel mechanism underlying systolic dysfunction in arrhythmogenic cardiomyopathy driven by a desmoplakin mutation that causes functional haploinsufficiency. Leveraging both 3D and 2D iPSC-derived cardiomyocyte models, we faithfully recapitulate key features of the disease and reveal that altered force transmission at the intercalated disc disrupts the regulation of sarcomere length, a previously unrecognized contributor to the pathogenesis of this disorder.

### Resource Availability

#### Lead Contact

Requests for further information and resources should be directed to and will be fulfilled by the lead contact, Stuart Campbell

#### Materials Availability

This study did not generate new unique reagents.

#### Data and Code Availability

This study did not generate DNA or RNA sequencing results; proteomics; metabolomics; or structural data. Original and uncropped Western blot images have been included in Supplementary Fig. S8. Microscopy data will be shared by the lead author upon reasonable request.

## Acknowledgements

We are grateful to Dr. Brent Hoffman (Duke University) for insightful discussions on 2D immunostaining, and to Dr. Akira Nagafuchi (Nara Medical University) for generously sharing the α18-catenin antibody. Confocal microscopy images were obtained at the Yale Science Hill Imaging Core.

## Author Contributions

Conceptualization: S.G.C., F.G.A., and I.G.; methodology: S.G.C. and I.G.; investigation: I.G., X.L., J.S., A.J.M.P.; writing (original draft): I.G.; writing (review): all authors; funding: S.G.C. and F.G.A.; supervision: S.G.C. Artificial intelligence tools were not used in the production of this manuscript.

